# The freeze-avoiding mountain pine beetle (*Dendroctonus ponderosae*) survives prolonged exposure to stressful cold by mitigating ionoregulatory collapse

**DOI:** 10.1101/2024.02.14.580279

**Authors:** Mads Kuhlmann Andersen, Amanda Diane Roe, Yuehong Liu, Antonia Musso, Serita Fudlosid, Fouzia Haider, Maya Evenden, Heath A. MacMillan

**Author notes:** Corresponding author: Mads Kuhlmann Andersen.

## Abstract

Insect performance is intrinsically linked to environmental temperature, and surviving through winter represents a key challenge for temperate, alpine, and polar species. To overwinter, insects have adapted a wide range of strategies to become truly cold hardy. However, while the physiological mechanisms underlying the ability to avoid or tolerate freezing have been well-studied, little attention has been given to the challenge of maintaining ion homeostasis at frigid temperatures in these species, despite this being a central issue for insects susceptible to mild chilling. Here we investigate how prolonged exposure to temperatures just above the supercooling point affects ion balance in freeze-avoiding larvae of the mountain pine beetle (*Dendroctonus ponderosae*) in autumn, mid-winter, and spring, and relate it to organismal recovery times and survival outcomes. We found that hemolymph ion balance was gradually disrupted during the first day of exposure, characterized by hyperkalemia and hyponatremia, after which a plateau was reached and maintained for the rest of the seven day experiment. The degree of ionoregulatory collapse experienced by larvae correlated strongly with recovery times, which followed a similar asymptotical progression. Mortality increased slightly during the most severe cold exposures, where hemolymph K^+^ concentration was highest, and a logistic relationship was found between survival and hyperkalemia. Thus, the cold tolerance of the freeze-avoiding larvae of *D. ponderosae* appears limited by the ability to prevent ionoregulatory collapse in a manner similar to less tolerant chill-susceptible insects, albeit at much lower temperatures. Furthermore, we posit that a prerequisite for the evolution of insect freeze avoidance is a convergent or ancestral ability to maintain ion homeostasis during exposure to extreme cold stress.

## Introduction

Most insects have limited capacity for thermoregulation, so their body temperature closely matches that of the environment (Heinrich, 2013; Lahondère, 2023). Their physiology is therefore intimately tied to external temperature, making performance and survival in extreme or variable thermal environments a key challenge (Harrison et al., 2012). This is particularly true for temperate, alpine, and polar insect species which often need to survive through extended periods of sub-zero temperatures during winter, a period that can represent a large proportion of their life span (Sinclair et al., 2003; Williams et al., 2015). For these reasons, low temperature survival and overwintering success are continuously highlighted as key factors in determining insect abundance and distribution (Bale, 1987; Addo-Bediako et al., 2000; Ramløv, 2000; Bale, 2002; Marshall et al., 2020). This is also true for the mountain pine beetle, *Dendroctonus ponderosae*, a pest native to western North America, that has spread beyond its native range and invaded regions east of the Rocky Mountains that experience more severe winter conditions than its native range (Safranyik et al., 2010; de la Giroday et al., 2012; Creeden et al., 2014). Thus, exploring the cold tolerance physiology of *D. ponderosae* is critical to understanding the processes that promote or limit its continued invasion into the eastern boreal forest.

Insects have acquired a wide range of adaptations and strategies that allow them to adjust to, thrive in, or at the very least survive, low temperature environments. Insect cold tolerance strategies and cold hardiness have historically been divided into two main categories; strategies that do and do not involve temperatures low enough to cause freezing of the extracellular fluid (Lee, 1991; Bale, 1996; Denlinger and Lee Jr, 2010; Lee, 2012). The majority of insects belong to the latter group and are characterized as chill-susceptible, meaning they succumb to relatively mild cold at temperatures above those that cause extracellular freezing (Overgaard and MacMillan, 2017). For these insects, survival depends on overcoming the direct, disruptive effects of low temperature on homeostasis, cellular integrity, and molecular processes. Central to the current conceptual model of chill tolerance physiology is a cold-induced disruption of extracellular ion homeostasis, referred to as an ‘ionoregulatory collapse’ (MacMillan, 2019). Under benign conditions, the renal system (Malpighian tubules and hindgut) maintains hemolymph ionic and osmotic balance through a continuous cycle of secretion and reabsorption, which counteracts transepithelial and cellular leakage (Edney, 1977; Phillips, 1981; Dow, 2017). However, at low temperature the activity of ion-motive transporters responsible for secretion and reabsorption are slowed below the level required for maintenance of homeostasis (MacMillan and Sinclair, 2011; Overgaard et al., 2021). Thus, prolonged exposure to stressful cold leads to a debilitating increase in hemolymph K^+^ concentration (hyperkalemia), which, in combination with cold, depolarizes excitable tissues, triggers an intracellular calcium overload, and leads to cell death and organismal injury (MacMillan et al., 2015c; Bayley et al., 2018). Subsequently, recovery from cold exposure depends on the ability to either restore hemolymph ion concentrations upon return to permissive temperatures or prevent ionoregulatory collapse altogether (MacMillan et al., 2014). Within chill-susceptible insects, however, there is substantial variation in the amount of cold that species or populations can handle, which has prompted the establishment of a vaguely-defined separation of insects that are truly chill-susceptible from those that are chill tolerant, where the latter group is able to either prevent or mitigate the loss of ion homeostasis *via* a wide range of ionoregulatory adaptations and therefore tolerate slightly more severe cold exposures (Sinclair, 1999; Overgaard and MacMillan, 2017; Overgaard et al., 2021).

For other insects, cold survival is determined below or at temperatures that cause extracellular freezing. These species are divided into two main strategies: freeze tolerant and freeze avoidant (Bale, 1987; Bale, 1996). As the name suggests, freeze tolerant insects can survive freezing of the extracellular fluids and initiate freezing at relatively high sub-zero temperatures. This slows ice formation and help limit ice to the extracellular space (Salt, 1961; Zachariassen, 1985). Thus, these insects have a relatively high supercooling point (SCP), the temperature at which freezing occurs spontaneously during a cooling challenge (Sinclair, 1999; Sinclair et al., 2015). Freeze-avoiding insects, on the other hand, survive substantial cold by possessing the ability to supercool to extremely low temperatures (often lower than -30°C) (Sinclair et al., 2003). These species have adapted to survive through harsh winters without freezing by, for example, accumulation of cryoprotectants (i.e. compatible osmolites and anti-freeze proteins), removal of ice-nucleating agents, and cryoprotective dehydration (Salt, 1961; Sømme, 1982; Graham et al., 1997; Holmstrup et al., 2002). Each of these processes effectively lowers the SCP of the insects but are rarely found alone, meaning that the impressive supercooling capacity of freeze-avoiding insects is achieved *via* a combination of colligative non-colligative factors (Zachariassen, 1985; Block et al., 1990; Sinclair et al., 2003; Lee, 2012; Storey and Storey, 2012). For example, freeze-avoiding beetles accumulate low molecular weight polyols in preparation for overwintering and for producing a wide range of anti-freeze proteins which lower the limit for freezing by preventing the formation of nucleating ice crystals (Sømme, 1964; Storey and Storey, 1983; Gehrken, 1984; Graham et al., 2007).

*Dendroctonus ponderosae* typically overwinter as freeze avoiding larvae and possess a remarkable ability to lower their SCP in this life stage (Régnière and Bentz, 2007; Rosenberger et al., 2017). This reduction in SCP is mainly achieved through accumulation of substantial amounts of glycerol in response to emerging winter conditions (Bentz and Mullins, 1999; Robert et al., 2016; Thompson et al., 2020). This ability to survive severe cold is thought to contribute to its recent eastward spread of MPB from its native range across the Rocky Mountains, and into Alberta and northeastern British Columbia (Canada) (Safranyik et al., 1975; de la Giroday et al., 2012; Creeden et al., 2014), where it has devastated substantial areas of coniferous forest and required over $500M to manage its eastern spread (Cullingham et al., 2011; Hodge et al., 2017). However, despite an impressive capacity for freeze-avoidance, cold mortality remains a key factor in determining population abundance of *D. ponderosae* (Renault et al., 2002; Régnière and Bentz, 2007). Thus, *D. ponderosae* represent an interesting and valuable system to study the physiological mechanisms underlying pre-freeze mortality in freeze-avoiding insects, which are generally thought to survive unless frozen (Salt, 1961).

Little is known about the processes that limit winter survival of freeze-avoiding insects, independent of freezing, although everything from dysregulation of metabolism to disrupted development has been suggested (Bale, 1987; Bale, 1991; Sømme, 1999). For freeze-avoidance to be a viable cold tolerance strategy, its evolution must have included an ancestral or convergent ability to avoid ionoregulatory collapse. If this is the case, we expect that freeze avoiding species can maintain ion balance during exposure to very low temperatures just above the lethal SCP. If ion homeostasis is disrupted in a freeze-avoiding insect, however, this could indicate an unappreciated role for ionoregulatory collapse in limiting the cold survival of freeze-avoiding species, which could indicate that, much like chill-susceptible insects, variation in ionoregulatory capacity dictates cold survival. Interestingly, phytophagous bark beetles, like *D. ponderosae*, possess unique hemolymph ion compositions with very low concentrations of Na^+^ and elevated concentrations of K^+^, Mg^2+^, and Ca^2+^ (Duchăteau et al., 1953; Jeuniaux, 1971). In other insects, low extracellular Na^+^ is associated with increased cold tolerance via mitigation of ionoregulatory collapse (MacMillan et al., 2015b; Lebenzon et al., 2020). Transmembrane concentrations of Na^+^ can play a role in freeze-tolerance and -avoidance, however, the evidence for this remains scarce (Zachariassen et al., 2004). Thus, exploring ion balance and its regulation in the freeze-avoiding *D. ponderosae* larvae might provide insight into evolutionarily conserved cold tolerance mechanisms.

In the present study, we investigated whether ion balance plays a role in limiting the cold tolerance of the freeze-avoiding *D. ponderosae* and asked the following main question: Do *D. ponderosae* suffer from ionoregulatory collapse when exposed to prolonged periods of stressful cold immediately above those that cause extracellular freezing? If this extremely cold-hardy insect suffers from ionoregulatory collapse in the cold, renal mechanisms of failure may be critical to fully understanding overwintering success of this and other freeze avoiding species. If, on the other hand, *D. ponderosae* do not suffer from ionoregulatory collapse, mechanisms that serve to defend against this disruption in the cold represent undescribed means by which freeze avoiding insects acquire cold tolerance (Overgaard and MacMillan, 2017; MacMillan, 2019; Overgaard et al., 2021). Given that non-freezing cold mortality occurs during overwintering in this species (Régnière and Bentz, 2007), we hypothesized that this could be linked to a gradual loss of ion balance (i.e. hemolymph hyperkalemia). Here, we exposed *D. ponderosae* that were in 1) preparation for overwintering, 2) in the middle of overwintering, and 3) emerging from overwintering to extended periods of stressful cold (above their SCP). We obtained parallel measurements of cold tolerance phenotypes and hemolymph ion concentrations at select time points that yielded a dataset that demonstrates mountain pine beetles can partly, but not completely, mitigate ionoregulatory collapse in the cold during overwintering.

## Materials and Methods

### Animal collection sites and husbandry

All specimens of *Dendroctonus ponderosae* (Hopkins, 1902) used in this study were collected from a population from its expanded range in central Alberta. To collect experimental animals, adult D. ponderosae were baited to lodgepole pine trees (*Pinus contorta* var. *latifolia*, Engelmann) at a field site in the expanded range of *D. ponderosae* outside of Cynthia, Alberta (Canada; 53.34 N, - 115.48W; see **Fig. 1**). Baiting was achieved using four funnel traps containing aggregation pheromones (trans-verbenol and exo-brevicomin) and host tree volatiles (myrcene and terpinolene) (product #3093, Synergy Semiochemical, Delta, Canada V4G 1E9) for five weeks during July and August of 2022. Here, attracted beetles mass attacked the trees, which were then left on the landscape for egg laying and brood development until harvest. In October 2022, seven trees were harvested and cut into 50 bolts (∼ 40-50 cm long and ∼ 30-50 cm in diameter), and the ends were sealed with paraffin wax to prevent desiccation. Infested bolts were transported to the Department of Biological Sciences at University of Alberta for outdoor storage and overwintering. Temperature-loggers (HOBO 2x External Temperature Data Logger [Onset, Bourne, MA, USA]) were implanted underneath the bark of three separate logs to monitor the thermal environment of the larvae during the overwintering period. Under bark recordings could only be started after trees were harvested and bolts positioned for storage overwinter, so the prior thermal history of the trees was approximated by obtaining and averaging temperature data from three weather stations located near the field site (https://climate.weather.gc.ca/index_e.html, see **Fig. 1** and **Table S1** for locations). At three points during the overwintering period, logs were transported (by air, overnight, and in double-walled secure containers with export authorization from Alberta Agriculture, Forestry and Rural Economic Development, permit #UofA-03-2022) to the Insect Production and Quarantine Laboratory at the Great Lakes Forestry Centre (GLFC) in Sault Ste. Marie, ON (part of Natural Resources Canada, Canada P6A 2E5) where experiments on live animals were carried out in a PPC2A quarantine facility. The time points were chosen such that they included 1) larvae from mid-autumn at the very beginning of overwintering (October 25, 2022), 2) overwintering larvae from mid-winter (January 24, 2023), and larvae towards the end of overwintering in spring (April 12, 2023). Ten, 17, and 23 logs were transported to GLFC in October, January, and April, respectively, and after their arrival they were stored in the dark at 6°C until extraction for experiments (which was within nine days). Extraction was carried out at room temperature (21-22°C) and larvae (only 4^th^ instar) were used in experiments within an hour of being removed from the log.

**Fig. 1.**
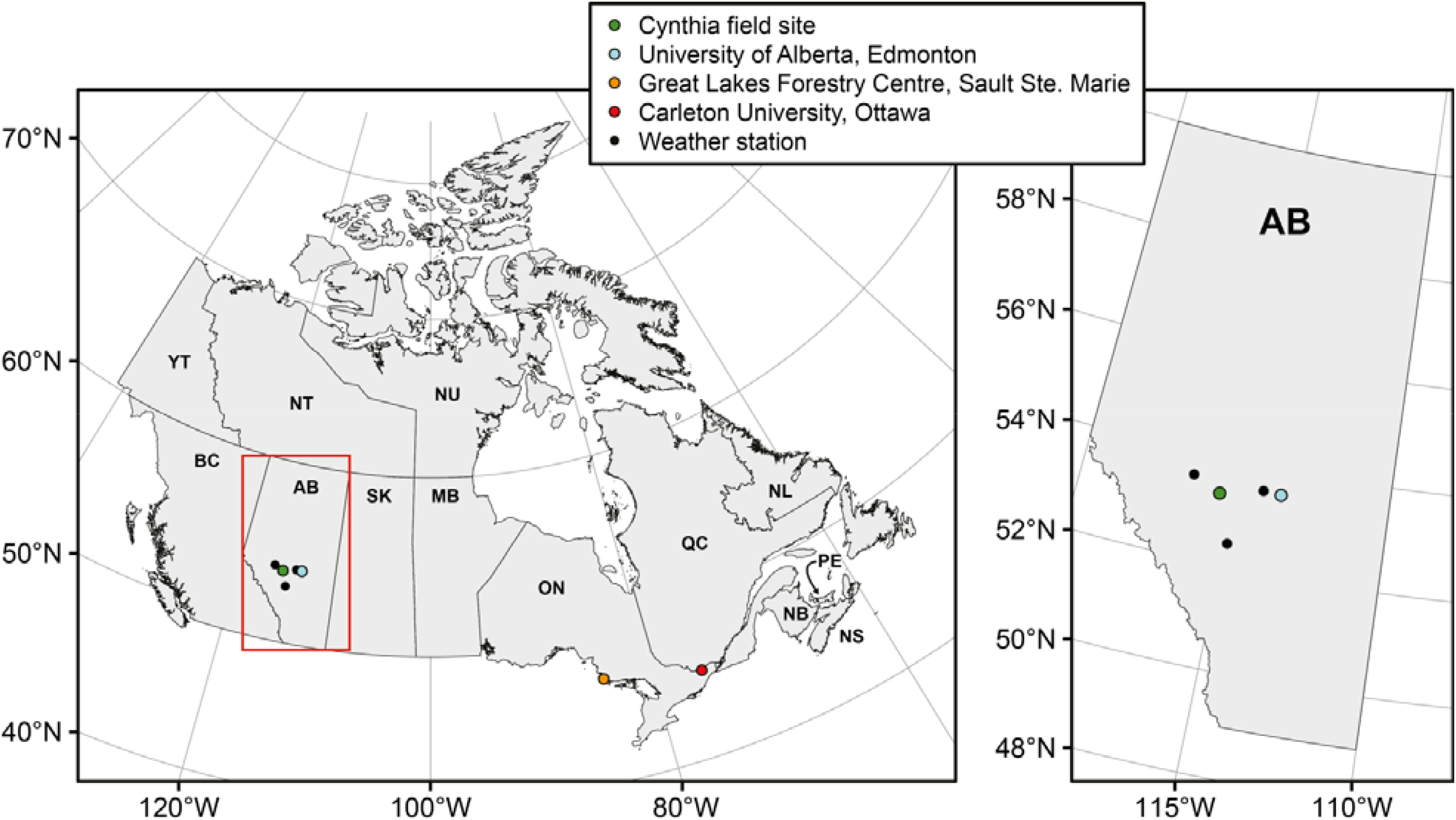
Map overview of the field site, and experimental locations. The experimental field site was just outside of Cynthia (Alberta, Canada T0E 0K0, green circle). Here, trees were baited, monitored for beetle attack, and finally harvested and brought to the Department of Biological Science at the University of Alberta (Edmonton, Alberta, Canada T6G 2E9, blue circle) for storage. At specific time points, subsamples of logs were transported to the quarantine facilities at the Great Lakes Forestry Centre (Natural Resources Canada, Sault Ste. Marie, Ontario, Canada P6A 2E5, orange circle) where experiments on live animals were carried out with permission. Hemolymph samples collected here were transported to the Department of Biology at Carleton University (Ottawa, Ontario, Canada K1S 5B6) where ion concentrations were measured. The bark temperature of the logs stored at University of Alberta was measured continuously, and temperature at the field site was approximated using hourly temperature measurements from three nearby field stations (smaller black circles) (see **Fig. 2** for temperature profiles).

### Cold tolerance measurements

At each of the three seasonal sampling points, we started by measuring supercooling points (SCP) of a subset of extracted *D. ponderosae* larvae before sampling larvae for all other experiments. SCPs were measured following the approach outlined by Sinclair et al. (2015). First, type-K thermocouples were gently attached to the outside of each larva with a small amount of vacuum grease, after which individual larvae were put into 200 μL microcentrifuge tubes. These were then gently stopped with cotton and placed into a custom-built aluminum casing through which a cooling bath (Proline RP855; Lauda, Würzburg, Germany) circulated a 1:1 water-methanol solution. From here, larvae were kept at 6°C for 10 min after which the temperature was lowered by 0.5°C min^-1^ until the temperature reached -35°C to ensure that all animals froze. At the end of the experiment, temperature traces were analyzed for each animal and the temperature at which the freezing exotherm occurred was recorded as the SCP. In this and subsequent experiments, all frozen animals were confirmed dead. A total of 53 larvae were used for this experiment with 19, 22, and 12 in autumn, winter, and spring, respectively.

After establishing the supercooling capacity of the larvae, we chose exposure temperatures that we expected to be stressful, but also minimized the risk of freezing. We chose 1) -10°C in the autumn, 2) -10°C and -20°C in the winter, and 3) -10°C in the spring, which represented 0.07 %, 0.01 %, 7.33 %, 7.59 % risk of reaching the SCP, respectively (based on standard deviations and assumed normal-distributed datasets). This approach does not account for the stochastic nature of freezing in cold-exposed insects (Ditrich, 2018), however, frozen larvae were easy to detect and were discarded (0, 0, and 17 larvae froze in the autumn, winter, and spring, respectively, representing 0, 0, and ∼ 6.9 % of larvae in these experiments). The -10°C exposure in the winter-acclimatized animals was included to allow for direct comparisons across seasons. From here, larvae were exposed to their respective temperature treatments by first holding them at 6°C for 10 min, before lowering the temperature by 0.5°C min^-1^ until -10°C or -20°C was reached and holding at that temperature for up to 168 h (seven days). During this cold exposure, larvae were removed from the cold at pre-determined time points at 0 h (immediately upon reaching the exposure temperature), 4 h, 16 h, 24 h, 48 h, 96 h, and 168 h.

At the specified time points, larvae were removed from the cold treatments and returned to room temperature (21-22°C) to estimate chill coma recovery time and survival as described for other insect larvae (Sinclair et al., 2015); any frozen larvae were discarded. Briefly, we removed larvae rapidly from tubes and placed them on a piece of paper at room temperature to observe and time the onset of movement for each larva. During this time, larvae were encouraged to move by gently poking them with a fine brush. Larvae were omitted from the data analyses if they did not move within 90 min. After the 90 min observation period, larvae were moved back into their tubes, which were gently stopped with cotton, and placed in the dark for 24 h at room temperature. Subsequently, their survival was estimated after the 24 h period by again placing them on a piece of paper and scoring them based on the ability to move as follows: 2 = spontaneous movements (alive, uninjured), 1 = only moving when stimulated (alive, injured), 0 = no movements (dead). A total of 390 larvae were used in these experiments (see Supplementary Data for distribution between seasons and treatments).

### Hemolymph collection and ion concentrations

In parallel with the experiment described above, larvae exposed to the same conditions were sampled for hemolymph after 0 h, 4 h, 16 h, 24 h, 48 h, 96 h, and 168 h after being held at either - 10°C or -20°C. Additional hemolymph samples were collected immediately after beetle extraction from the logs at room temperature (21-22°C) and after 24 h at 6°C following extraction. To sample hemolymph, we followed the procedure described for larvae by Campbell et al. (2018). Briefly, after removal from the cold treatments, larvae were rapidly submerged in hydrated paraffin oil in a Petri dish. A 26G needle or a pair of micro scissors was used to puncture the dorsal cuticle of the pro- or metathorax. The extruding hemolymph droplets from individual larvae were collected with a micropipette, transferred to a round-bottomed 96-well PCR plate, and stored under hydrated paraffin oil at -80°C. Hemolymph samples were shipped to the Department of Biology at Carleton University in Ottawa on dry ice and stored at -80°C until hemolymph ion concentrations were measured. A total of 413 hemolymph samples were collected in these experiments (see Supplementary Data for distribution between seasons and treatments).

On the day of ion measurement, hemolymph samples were thawed on ice and transferred to a glass Petri dish (60 mm diameter) with an elastomer covering the bottom (Sylgard 184, Dow Corning Corporation, Midland, MI, USA) containing hydrated paraffin oil. Here, the concentrations of Na^+^ and K^+^ were measured using ion-selective microelectrodes as described in MacMillan et al. (2015a). The Na^+^ concentration was measured only at a subset of time points. Ion-selective electrodes were fashioned from borosilicate glass (TW150-4; World Precision Instruments, Sarasota, FL, USA), pulled on a Flaming-Brown P-1000 micropipette puller (Sutter Instruments, Novato, CA, USA) to a tip diameter of ∼ 1-3 μm, and silanized at 300°C in an atmosphere of N,N-dimethyltrimethyl silylamine (Sigma Aldrich, St. Louis, MO, USA). From here, electrodes were back-filled with 100 mM KCl or NaCl depending on which ion was being measured and front-filled with the associated ionophore. For K^+^ measurements we used K^+^ ionophore I, cocktail B (Sigma) and for Na^+^ measurements we used a Na^+^ ionophore X (Sigma) cocktail designed to optimize Na^+^ selectivity (Messerli et al., 2008). Once filled, electrode tips were dipped in a solution of polyvinyl-chloride in tetrahydrofuran (10 mg in 3 mL; Sigma) to prevent ionophore displacement. Raw voltage outputs from the ion-selective electrodes were connected to a pH Amp (ADInstruments, Colorado Springs, CO, USA), digitized using a PowerLab 4SP A/D converter (ADInstruments), and read by a computer running Lab Chart 4 software (ADInstruments). The circuit was completed with a thinly pulled borosilicate glass electrode (1B200F-4, WPI) filled with 500 mM KCl. Voltages were converted to ion concentrations by comparison to standard with a ten-fold difference in concentration of the target ion (K^+^: 10 mM and 100 mM, Na^+^: 2.5 mM and 25 mM; difference made up with LiCl) and the following formula:

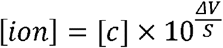

where [*ion*] is the concentration of the target ion, [*c*] is the ion concentration in one of the standards, *ΔV* is the difference in voltage between the sample and standard, and *S* is the difference in voltage between the two standards. The latter voltage response should be close to Nernstian (58.2 mV per ten-fold change in concentration) and was 55.1 ± 1.7 mV (N = 46) and 55.6 ± 1.7 mV (N = 37) for K^+^ and Na^+^ electrodes (means ± s.d.), respectively.

### Statistical analyses

All data analyses was performed in R software (v4.2.2, R Core Team (2023)). Data normality was tested with Shapiro Wilks tests, and homoscedasticity was tested for normally distributed datasets with F tests. Supercooling points were compared between seasonal sampling points with a Kruskal-Wallis rank sum test followed by pairwise Mann-Whitney-Wilcoxon tests with a Benjamini-Hochberg correction. The effects of season, exposure time, and exposure temperature on chill coma recovery times and survival were analyzed using linear models. Concentrations of Na^+^ and K^+^ were first compared with control groups (24 h at 22°C, 24 h at 6°C, and 0 h at each exposure temperature) with one-way ANOVAs. These analyses found no differences, so the effects of season, exposure time, and exposure temperature were analyzed with linear models while excluding the room temperature and 6°C controls for each ion. The relationships between chill coma recovery time, survival, and hemolymph K^+^ concentration were all analyzed with linear regression, linear regression on log-transformed data (i.e. an exponential relationship), or non-linear regression to a logistic model using the nls() function. The best fitting models were chosen based on AIC scores. For all analyses, the level of statistical significance was 0.05, and values reported represent means and their standard error (s.e.m.) unless stated otherwise. For sample sizes, see the Supplementary Materials (Tables S2-4).

## Results

### Overwintering conditions and cold tolerance of Dendroctonus ponderosae

To simulate overwintering, tree logs containing *D. ponderosae* larvae were kept in an outdoor enclosure at the Department of Biological Sciences at University of Alberta. Here they experienced a natural winter and a range of temperatures from ∼ 10°C in mid-autumn to a minimum temperature of -26.3°C in mid-late December and a gradual climb back to ∼ 20°C towards mid-April with several cold snaps in-between (**Fig. 2A**). During this time, larval SCPs changed significantly (*X*_*2*_ = 40.4, df = 2, P < 0.001) from -15.7 ± 0.4°C in the autumn (**Fig. 2B**), down to -26.7 ± 1.0°C in mid-winter (**Fig. 2C**), and back to -12.6 ± 0.5°C in the spring (**Fig. 2D**).

**Fig. 2.**
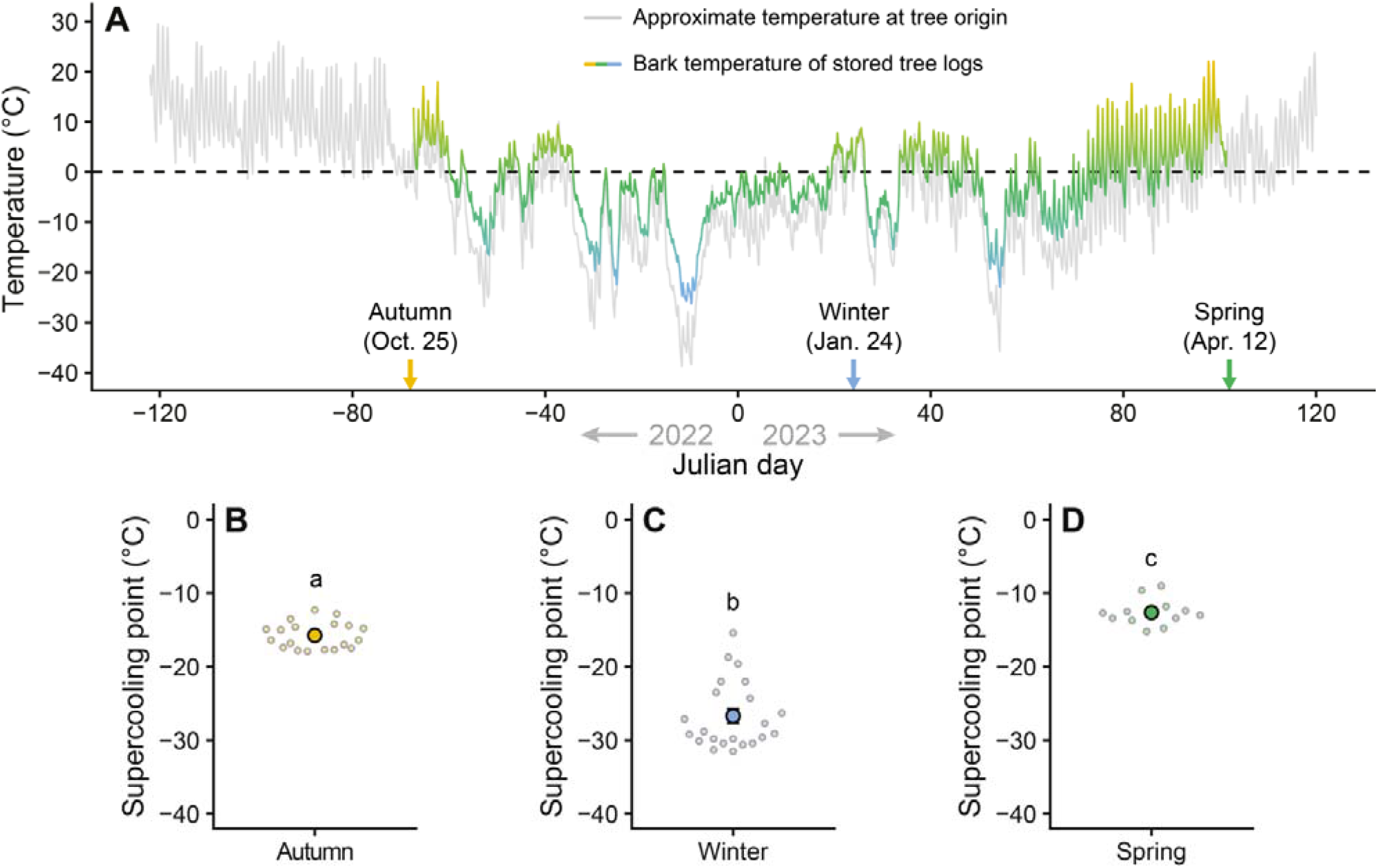
Temperature profiles and supercooling capacities of overwintering *Dendroctonus ponderosae* larvae from late 2022 to early 2023. During the overwintering of *D. ponderosae* from the autumn of 2022 to spring of 2023, temperature was tracked at their collection site using historical climate data from Environment Canada (**A**, grey), and monitored directly under the bark of logs that were moved into outdoor storage for experiments (**A**, coloured). Note that the trees that were cut into logs and stored for subsequent larvae extraction and experiments appear to have been sheltered from the most severe winter conditions (i.e. the coloured line is always above the grey line). At three time points during this period, larvae were extracted from the logs to measure their supercooling points (**B-D**). Small, translucent points denote individual data points, while the large opaque point denotes the mean. Error bars not visible are obscured by the symbols, and groups not sharing a letter are statistically different. For sample sizes, see Table S2.

At the same time, we estimated their ability to tolerate and survive prolonged exposure to extreme cold by measuring the chill coma recovery times and subsequent survival scores after each exposure (**Fig. 3**). The exposure temperatures were designed to be stressful while minimizing the risk of freezing, however, a few animals did freeze during the experiments performed in the spring, and these were removed from later analyses. We found that the recovery times increased with time exposed to stressful cold (F_1,357_ = 616.2, P < 0.001), decreasing temperatures (estimated only in winter; F_1,357_ = 4.7, P = 0.031), and their interaction (F_1,357_ = 7.6, P = 0.006) (**Fig. 3A-C**). At the same time, we found a clear effect of season on recovery times (i.e. autumn vs. winter vs. spring; F_2,358_ = 14.2, P < 0.001), which also shared a significant interaction with exposure times (F_2,357_ = 45.5, P < 0.001). Specifically, after having just reached their exposure temperatures at the end of the ramp down (i.e. the 0 h exposure), the recovery time was similar between all larvae exposed to -10°C (2.2 ± 0.1 min, 1.9 ± 0.3 min, and 2.4 ± 0.1 min, for autumn-, winter-, and spring-acclimatized larvae in orange, blue, and green circles, respectively) with winter-acclimatized larvae exposed to -20°C taking slightly longer (4.8 ± 0.8 min, blue triangles). From here, recovery times increased with exposure time and more-so at lower temperatures (measured in winter) and in the spring such that after 168 h these values had grown to 26.7 ± 3.5 min, 24.2 ± 3.8 min, 51.3 ± 2.7 min for -10°C exposed autumn-, winter-, and spring-acclimatized larvae, respectively, and to 44.1 ± 5.9 for winter-acclimatized larvae exposed to -20°C.

**Fig. 3.**
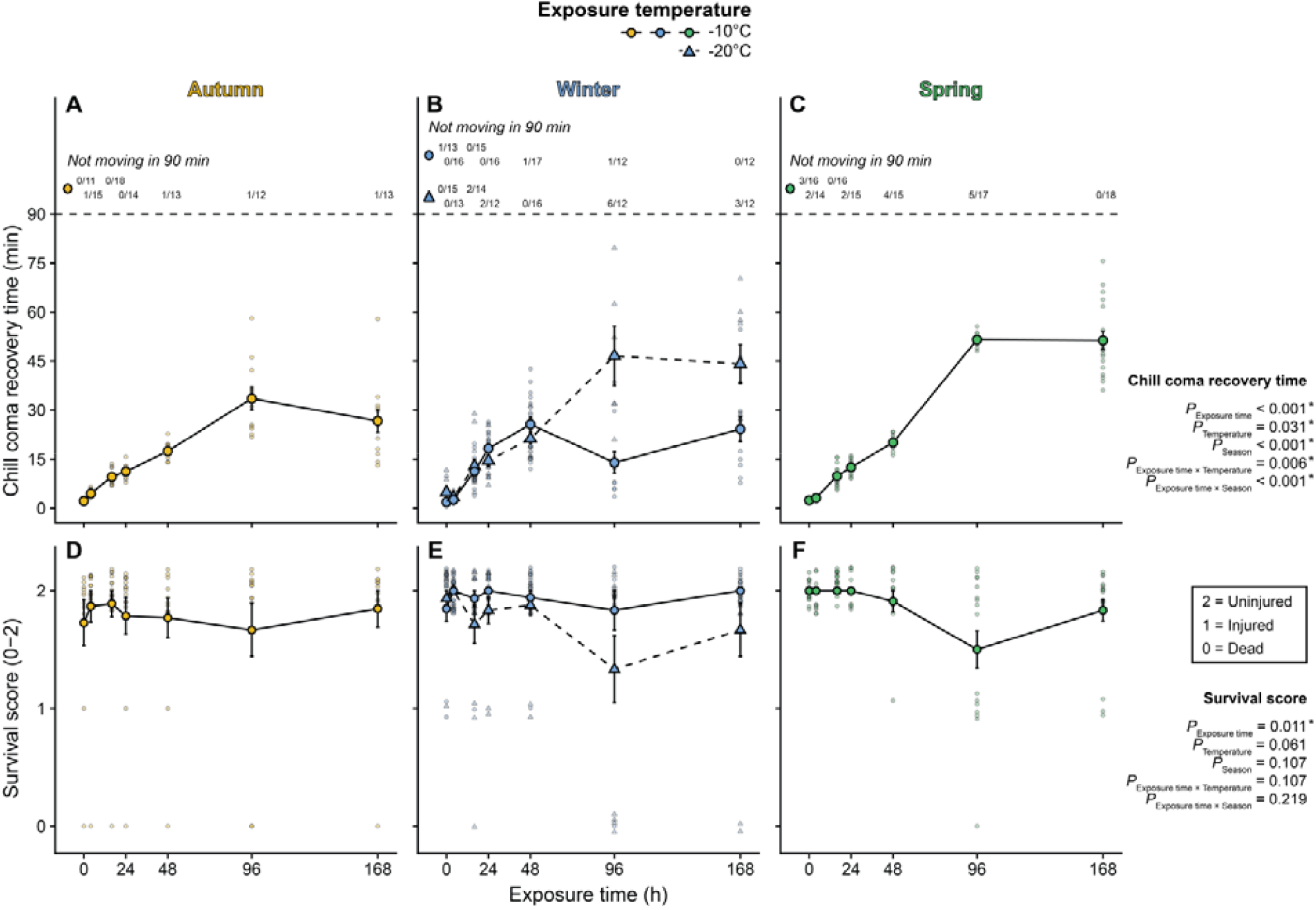
Cold tolerance measures of overwintering *Dendroctonus ponderosae* larvae. The cold tolerance of *D. ponderosae* larvae was estimated at three time points during the overwintering period from autumn of 2022 to spring of 2023 (see **Fig. 1** for dates). This was done by exposing them to periods of a stressfully low, non-freezing, coma-inducing temperatures (−10°C in circles with a full line, and -20°C in winter in triangles with a dashed line) up to 168 h (seven days) and estimating how long it took them to recovery mobility (**A-C**) and whether or not they accrued cold-related injuries (see Materials and Methods, **D-F**). Note that some larvae did not recovery function within the 90 min observation period (horizontal dashed line) and were removed from this dataset (**A-C**), however, unless frozen by the exposure, they were kept to assess their survival after the full recovery period. Overall, recovery times increased the longer larvae were exposed to cold stress; however, this was further prolonged if the exposure temperature was lowered or if the larvae had exited their overwintering stage (i.e. in spring). Survival, on the other hand, was only slightly decreased by extended periods of cold stress and was similar across seasons and temperature exposures. Small, translucent points denote individual data points, while the large opaque point denotes the mean. Individual data points on **panels D-F** are jittered along the ordinal y-axis for clarity. Error bars not visible are obscured by the symbols. For sample sizes, see Table S3.

After recovery from the cold exposure, larvae were left to recover at room temperature for 24 h, after which their survival score were estimated based on their ability to move. We saw very little evidence of injury in the larvae left to recover at room temperature for 24 h (**Fig. 3D-F**). Nonetheless, we identified a significant effect of exposure time on the survival score, as longer exposures slightly reduced the survival score (F_1,382_ = 6.6, P = 0.011). In addition, there was a marginally significant trend for exposure to lower temperatures to further reduce survival (F_1,382_ = 3.5, P = 0.061). Seasonality did not affect survival (F_2,382_ = 2.2, P = 0.107), and there were no interactions between exposure time and other fixed factors (Exposure time × Temperature: F_1,382_ = 2.6, P = 0.107; Exposure time × Season: F_2,382_ = 1.5, P = 0.219).

### Hemolymph ion composition during prolonged non-freezing cold exposure

To investigate whether the observed increase in chill coma recovery time and reduction in survival score could be the result of an ionoregulatory collapse, as seen in other insects, we took hemolymph samples from animals exposed to the same time and temperature treatments and measured the concentrations of K^+^ and Na^+^ with ion-sensitive electrodes (**Fig. 4**).

**Fig. 4.**
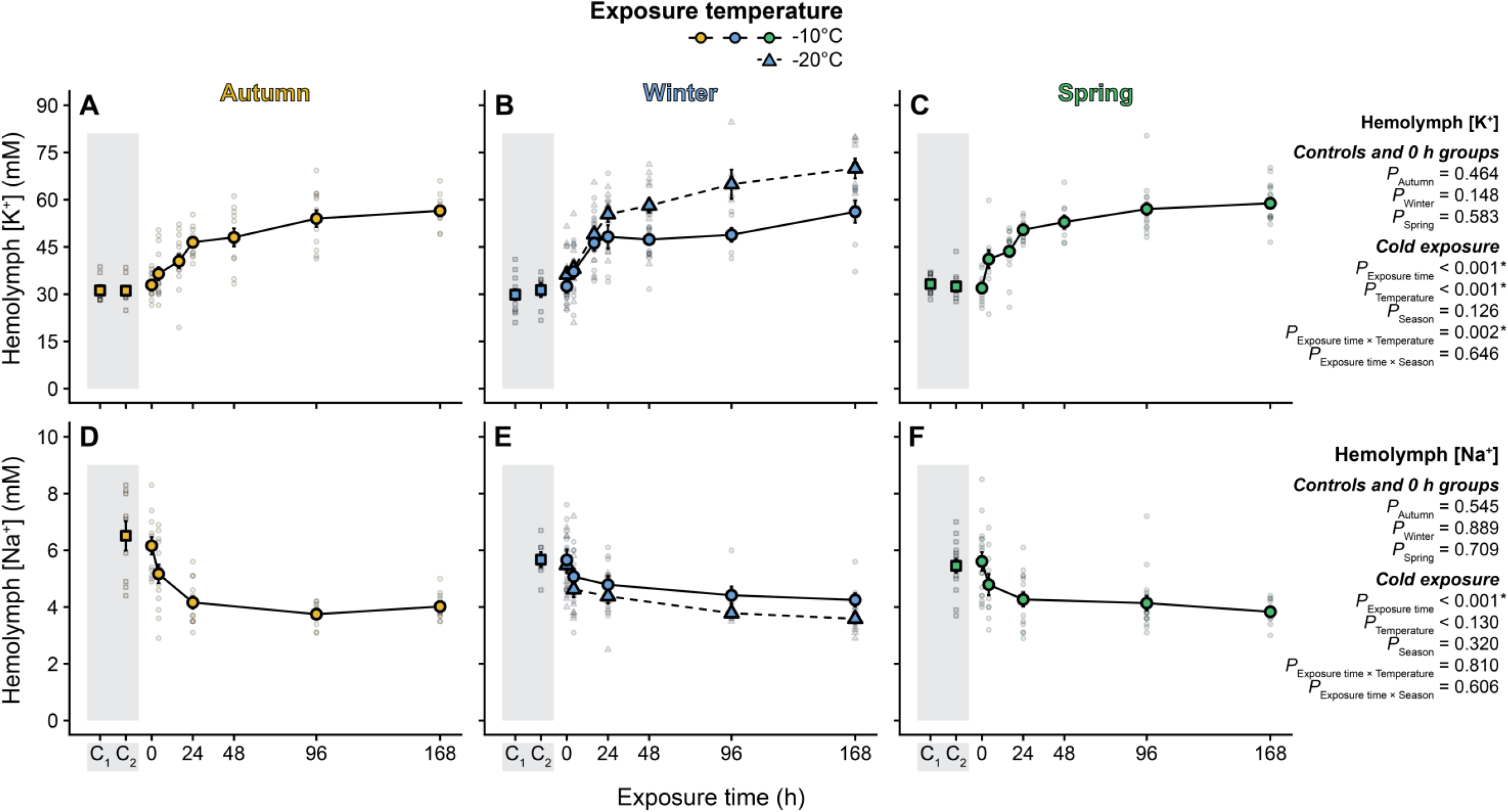
Prolonged exposure to stressful, non-freezing cold leads to moderate hemolymph hyperkalemia and hyponatremia in overwintering *Dendroctonus ponderosae*. After establishing that concentrations of K^+^ and Na^+^ in the hemolymph of larval *D. ponderosae* were unaffected be exposure to benign temperatures (squares in shaded area; C_1_ = 24 h at 21-22°C and C_2_ = 24 h at 6°C), larvae were again exposed to stressful cold (−10°C in circles with a full line, and -20°C in winter in triangles with a dashed line) for up to 168 h during which hemolymph was sampled. During the exposure hemolymph [K^+^] gradually increased, until an apparent asymptote was reached after a ∼ 80-82% increase above the control concentration in larvae held at -10°C (**A-C**). In -20°C-exposed larvae (winter), hemolymph [K^+^] increased further to ∼ 123% above baseline. Conversely, hemolymph [Na^+^] decreased gradually and generally reached concentrations ∼ 30-38% below baseline regardless of exposure temperature (**D-F**). Small, translucent points denote individual data points, while the large opaque point denotes the mean. Error bars not visible are obscured by the symbols. For sample sizes, see Table S4.

At each seasonal sampling point (autumn, winter, and spring) we first compared the hemolymph ion concentration in samples from control animals that were kept at room temperature (21-22°C), at 6°C for 24 h, at immediately upon reaching the exposure temperature (either -10 or - 20°C for 0 h). For Na^+^, however, we did not include the room temperature control. For hemolymph K^+^ concentration the values never differed between control groups (autumn: F_2,30_ = 0.8 and P = 0.464, winter: F_2,37_ = 1.9 and P = 0.148, spring: F_2,43_ = 0.5 and P = 0.583) and always stayed at ∼ 32 mM across seasons (mean ± s.d. = 32.3 ± 5.0 mM). The same was true for Na^+^ concentrations (autumn: F_1,19_ = 0.4 and P = 0.545, winter: F_2,26_ = 0.1 and P = 0.889, spring: F_1,29_ = 0.1 and P = 0.709) where the concentration stayed constant at ∼ 6 mM (mean ± s.d. = 5.8 ± 1.1 mM).

After confirming that our control conditions and very short cold exposure resulted in no change to hemolymph K^+^ and Na^+^ concentrations, we tested whether these were altered by prolonged exposure to stressful, non-freezing cold at -10°C, and at -20°C in winter. Hemolymph K^+^ concentration (**Fig. 4A-C**) increased with exposure time (F_1,335_ = 257.0, P < 0.001), with a lower exposure temperature (F_1,335_ = 35.4, P < 0.001), and with their interaction (F_1,335_ = 9.7, P = 0.002), with no effect of seasonality (F_2,335_ = 2.1, P = 0.126) nor interaction between seasonality and exposure time (F_2,335_ = 0.4, P = 0.646). Specifically, after 168 h at -10°C it increased from the control value of ∼ 32 mM to 56.5 ± 1.5 mM, 56.2 ± 3.5 mM, and 58.9 ± 1.5 mM in the autumn, winter, and spring, respectively, in a seemingly asymptotical manner, while it increased further to 69.9 ± 3.2 mM in the winter group exposed to -20°C. Conversely, the hemolymph Na^+^ concentration (**Fig. 4D-F**) decreased as exposure time increased (F_1,219_ = 63.3, P < 0.001) with no effects of exposure temperature (F_1,219_ = 2.3, P = 0.130), season (F_2,219_ = 1.1, P = 0.320), nor any interactions (time × temperature: F_1,219_ = 0.1 and P = 0.810, time × season: F_2,219_ = 0.5, P = 0.606). Specifically, over the time course of 168 h it decreased from the control value of ∼ 6 mM and stabilized at 4.0 ± 0.1 mM, 4.3 ± 0.2 mM, and 3.8 ± 0.1 mM, in the autumn, winter, and spring, respectively, and 3.6 ± 0.2 in the winter group exposed to -20°C.

### Correlations between hemolymph K^+^ and cold tolerance measures

Because we collected measurements of cold tolerance and hemolymph ion concentrations at the same time points throughout seasons and exposures, we can investigate the putative relationship between these parameters based on group means to investigate if this matches patterns observed for less cold tolerant, chill-susceptible insects (**Fig. 5**).

**Figure 5.**
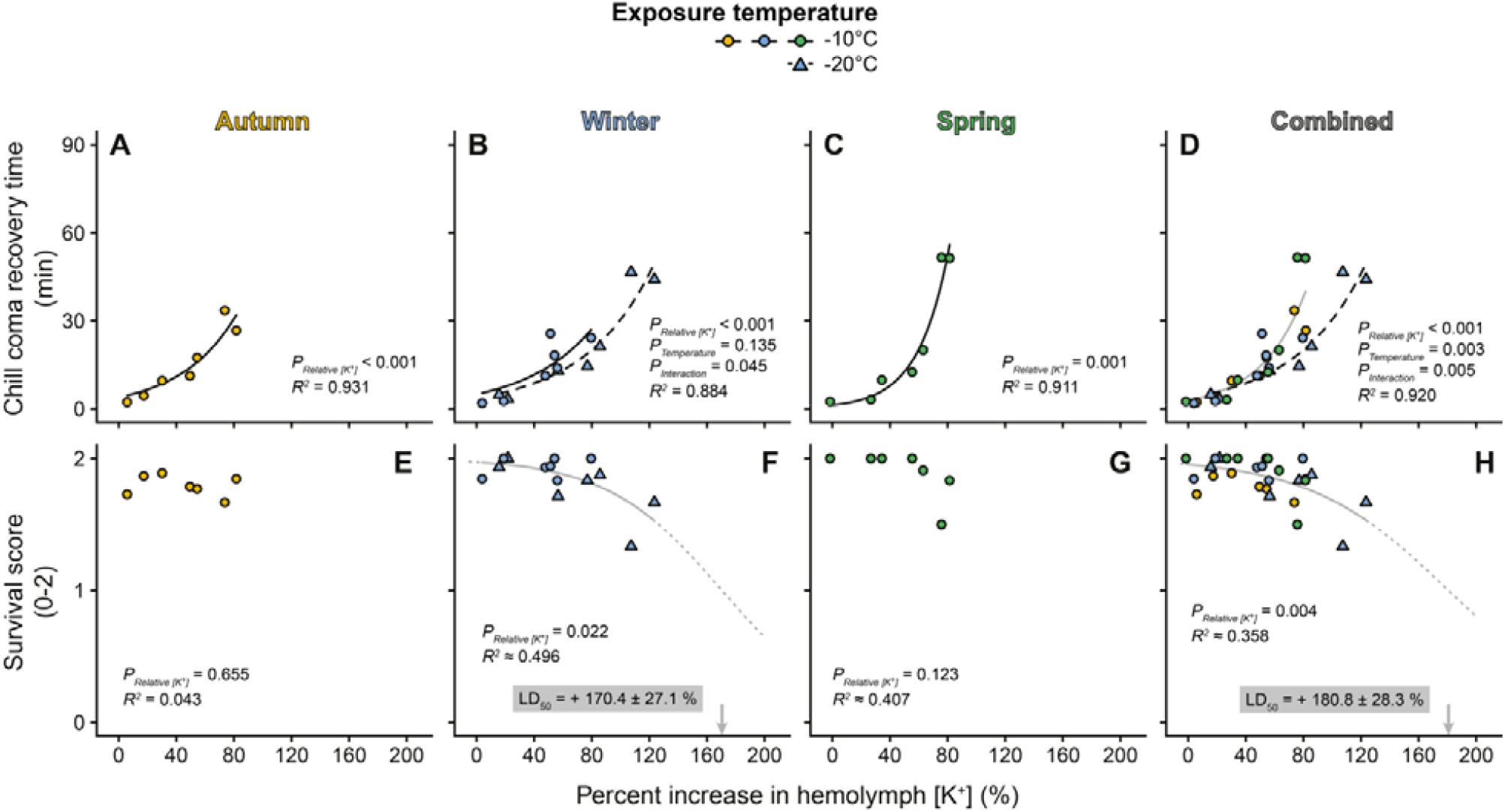
Magnitude of hemolymph hyperkalemia correlates with chill coma recovery times and survival outcome in cold-exposed, overwintering *Dendroctonus ponderosae* larvae. Correlations were made using the matching hemolymph [K^+^] and cold tolerance measurements at each time points across seasons and temperature exposures. Overall there was a strong exponential relationship between hemolymph hyperkalemia and the ability to recover movement after cold exposure (chill coma recovery time, **A-D**), and this relationship was right-shifted in larvae exposed to -20°C in mid-winter (**B** and **D**), indicating a faster rate of recovery at a similar degree of hyperkalemia. Exposure to -10°C for extended periods of time did not lead to any significant injury in the autumn (**E**) or winter (**F**). However, if larvae exposed to -20°C in winter were combined with those exposed to -10°C, the beginning of a sigmoidal dose-response curve emerged (**F**), and predicted that that the average larvae would be expected to have accrued injury at a 170.4 ± 27.1% increase in hemolymph [K^+^] (LD_50_, based on extrapolation). In spring, the relationship between survival score and hemolymph [K^+^] revealed a similar, yet statistically non-significant, sigmoidal curve with an LD_50_ of 110.3 ± 28.5% above control (**G**). With all datasets combined, we saw a significant relationship between hemolymph hyperkalemia and the decrease in survival score during the exposure to stressful cold (**H**) which also appeared sigmoidal and with a LD50 at a 180.8 ± 28.3 % increase hemolymph [K^+^]. Full lines and dashed black lines denote a significant relationship between the cold tolerance measure and the proportional increase in hemolymph [K^+^]; no line indicates the absence of a statistically significant relationship. A grey line denotes that the relationship is based on multiple treatment or seasonal groups (e.g. -10°C exposed groups across all seasons in **panel D**), while a dotted line represents a model-based extrapolation (in **panels F-H**).

We found a strong link between chill coma recovery time and hemolymph hyperkalemia, which closely matched an exponential relationship in the autumn (F_1,5_ = 67.8, P < 0.001), winter (F_1,10_ = 94.0, P < 0.001), and spring (F_1,5_ = 57.9, P < 0.001), as well as throughout the overwintering season when combined into one dataset (F_1,24_ = 227.6, P < 0.001) (**Fig. 5A-D**). Interestingly, we also found that a lower exposure temperature (in mid-winter) resulted in improved recovery rates, but only at higher degrees of hyperkalemia (**Fig. 5B**; hyperkalemia × temperature interaction: F_1,10_ = 5.2, P = 0.045) (effect of temperature: F_1,24_ = 2.6, P = 0.135). When datasets were combined, the faster recovery rate after exposure to lower temperature became statistically significant (F_1,24_ = 10.8, P = 0.003), and the effect remained larger at more severe hyperkalemia (F_1,24_ = 9.7, P = 0.005) (**Fig. 4D**).

The link between hemolymph hyperkalemia during cold exposure and survival outcome after recovery was less clear (**Fig. 5E-H**). In the autumn and spring, there were no apparent relationships (autumn, **Fig. 5E**: F_1,5_ = 0.2, P = 0.655; spring, **Fig. 5G:** F_1,5_ = 3.4, P = 0.123), however, a significant relationship emerged in winter (**Fig. 5F**; F_1,12_ = 6.9, P = 0.022). When combining data across seasons, the overall relationship between hemolymph hyperkalemia and survival was supported (**Fig. 5H**, F_1,26_ = 9.8, P = 0.004). The sigmoidal regressions fit to the statistical significant relationships showed that the degree of hemolymph hyperkalemia expected to injure all animals was an increase of ∼ 170-180% above control (i.e. ∼ 86-90 mM, **Fig. 4F**,**H**).

Similar significant relationships occurred between cold tolerance measures and hemolymph hyponatremia (**Fig. S1**). Specifically, we found an exponential relationship between chill coma recovery and hyponatremia (F_1,16_ = 53.0, P < 0.001), which was season-specific (effect of season: F_2,16_ = 5.1, P = 0.020), and a sigmoidal, dose-response-like relationship with the survival score (F_1,18_ = 5.4, P = 0.031) which indicated an onset of injury at a 72.0 ± 19.2 % decrease in hemolymph Na^+^ concentration.

## Discussion

In the present study, we examined how the cold tolerance of *D. ponderosae* larvae changed during overwintering and how potential non-freezing mortality might relate to ion balance disruption as seen in less tolerant insects. We show that overwintering larvae exposed to extremely low temperatures, just above those that cause them to spontaneously freeze, for prolonged periods experience physiological stress and sustain injury which manifested as increased recovery times and reduced survival. Stressed larvae experience a loss of ion balance, characterized by hyperkalemia and hyponatremia, which correlates strongly with the degree of cold-induced injury.

### Survival of overwintering mountain pine beetle larvae is only compromised after prolonged exposure to severe non-freezing cold

Previous studies demonstrated that overwintering larvae of *D. ponderosae* are freeze avoiding with the ability to substantially lower their SCP during winter (Régnière and Bentz, 2007; Rosenberger et al., 2017). Here we reaffirm that overwintering larvae in the expanded range are able to lower their supercooling points, and also show that dysregulation of ion balance and non-freezing chilling injury occur during prolonged chilling at ecologically relevant temperatures in this species.

The SCP we observed in mid-winter were well below the lowest observed temperature of the stored logs (see **Fig. 2**). This extensive SCP depression is a defining characteristic of freeze-avoiding species (Zachariassen, 1985), and the lowest SCP observed here is similar to that found in another beetle, the emerald ash borer (*Agrilus planipennis*, see Crosthwaite et al. (2011) and Duell et al. (2022)). During our experiments we noted that fall-collected larvae had food in their guts, while those extracted in winter and spring had empty guts. This is a common strategy for freeze-avoiding insects, as this removes most internal ice nucleating agents. We also noted some fall larvae held for prolonged periods at-10°C evacuated their guts during the experiment.

Mechanisms that suppress the SCP, and thereby reduce the risk of freezing, have historically been considered as the primary means by which freeze avoidant insects survive winter. The potential for non-freezing injury has received considerably less attention in freeze avoidant species. For less cold tolerant (chill-susceptible) insects, survival is determined by physiological effects of low temperature in the absence of freezing. In these species, low temperature tolerance is estimated by assessing the capacity to recover from and survive a non-freezing cold exposure (Sinclair et al., 2015; Overgaard and MacMillan, 2017). Our findings reveal that larvae of *D. ponderosae* exposed to prolonged periods of low temperatures just above their SCP were cold-stressed; recovery times gradually increased with exposure time and survival decreased slightly after prolonged exposure. These effects of cold exposure were exacerbated at lower temperatures and during seasons of reduced cold tolerance (i.e. spring, see **Fig. 3**). These trends are remarkably similar to those observed for chill-susceptible insects where both cold acclimation and milder exposure temperatures improve recovery times and survival outcomes (Andersen et al., 2017a; Yerushalmi et al., 2018; Tarapacki et al., 2021). Similarly, recovery times in other chill-susceptible insects reach an apparent plateau as injury and mortality begins to set in (MacMillan et al., 2012). That being said, the progression of adverse effects in *D. ponderosae* is markedly slower than that for chill-susceptible insects, even though the larvae were exposed to temperatures that would rapidly kill less tolerant insects (Koštál et al., 2004; Koštál et al., 2006; Findsen et al., 2013; Andersen et al., 2015). Interestingly, chill tolerant *Drosophila montana* recover rapidly following prolonged cold stress without injury, which could suggest shared physiological mechanisms to survive low temperatures (MacMillan et al., 2015a; Andersen and Overgaard, 2019). However, despite *D. ponderosae* larvae being able to mitigate the cascade of events that prolongs recovery times, they eventually suffer injury and mortality, indicating that they are unable to completely counteract the negative effects of the cold stress.

### Ion balance and partial ionoregulatory collapse during stressful cold exposure in mountain pine beetle larvae

In the ‘ionoregulatory collapse’ model of insect cold tolerance, increased recovery times and decreased survival are linked to a progressive, debilitating cold-induced hemolymph hyperkalemia (Overgaard and MacMillan, 2017; MacMillan, 2019; Overgaard et al., 2021). The hemolymph composition of phytophagous coleopterans has previously been described as being “unconventional”; they often have high concentrations of K^+^ (> 30 mM) and Mg^2+^ (> 30 mM), and unusually low concentrations of Na^+^ (< 15 mM) (Duchăteau et al., 1953; Jeuniaux, 1971). This is also the case in *D. ponderosae*. Hemolymph concentrations of K^+^ and Na^+^ were ∼ 32 mM and ∼ 6 mM, respectively, for *D. ponderosae* larvae kept under benign conditions (Mg^2+^ was not measured) (**Fig. 4**). As in chill-susceptible species, these values remained unchanged during the cooling process (MacMillan et al., 2014), but prolonged exposure caused hemolymph hyperkalemia and hyponatremia. Further, the degree of change in the concentration of these ions strongly correlated with the severity of the cold exposure and relative cold tolerance of the larvae (compare **Fig. 3** and **4**). Cold-induced hemolymph hyperkalemia is a hallmark of the ionoregulatory collapse model derived mainly from the study of dipteran and orthopteran species that are chill-susceptible (Overgaard and MacMillan, 2017; Overgaard et al., 2021). The same phenomenon has occurred in a chill-susceptible coleopteran, *Alphitobius diaperinus* (Koštál et al., 2007), and chill-susceptible lepidopterans with an ion homeostasis similar to that of *D. ponderosae* (Natochin and Parnova, 1987; Andersen et al., 2017b). Thus, despite being categorized as a freeze-avoidant insect (Régnière and Bentz, 2007; Rosenberger et al., 2017), *D. ponderosae* appears to be limited in its non-freezing cold tolerance via mechanisms that closely resemble those limiting chill-susceptible insects. Indeed, when the degree of hemolymph hyperkalemia correlates with recovery times and survival outcomes at the different time points, the emerging relationships resemble those found for chill-susceptible insects (**Fig. 5**). Specifically, there is a strong exponential correlation between the increase in hemolymph [K^+^] and recovery times, while the relationship with survival appears sigmoidal in nature (yet we caution that the fitted sigmoidal relationships extrapolate beyond the data). In chill-susceptible insects, the exponential relationship between recovery times and hemolymph hyperkalemia has been suggested to stem from the need for water-redistribution after the cold-induced osmotic disruption (MacMillan et al., 2015a; MacMillan et al., 2015b). In the present study, we did not investigate water balance, however, the cold-induced osmotic disruption in fruit flies is caused by a steep hemolymph-to-gut Na^+^ gradient which is unlikely to exist in *D. ponderosae* larvae as control concentrations are ∼ 6 mM. Nonetheless, hemolymph hyperkalemia does occur, and the approximate LD_50_ found here (∼ 170-180 % increase, see **Fig. 5**) closely matches that for chill-susceptible insects (∼ 200% increase, see Overgaard and MacMillan (2017)). Thus, even if different upstream homeostatic disruptions cause hyperkalemia in this species, the mechanism(s) by which increased hemolymph [K^+^] causes injury and cell death during a cold exposure might be shared.

### A distinct pattern of ion balance disruption in mountain pine beetles leads to new paths of inquiry on the mechanisms of seasonal plasticity and cold tolerance evolution

Intriguingly, the level of hemolymph hyperkalemia we observed in *D. ponderosae* plateaus at a level just below those that start to cause extensive mortality in other chill-susceptible species (i.e. they stabilize below a ∼ 150% increase while a ∼ 200% increase leads to injury, see Overgaard and MacMillan (2017)). This suggests that this freeze avoidant species has previously undescribed mechanisms that help prevent extensive cell depolarization, cell death, and organismal mortality in the cold. These mechanisms could already be established through seasonal plasticity before cold stress (i.e. seasonal plasticity or acclimatization), or could be induced rapidly during cold stress (i.e. rapid hardening), or a combination of both.

In chill tolerant *Drosophila* species or cold acclimated *D. melanogaster*, concentration gradients for the prevailing cation in the hemolymph (Na^+^) are reduced, and other compatible osmolytes are more abundant and serve to reduce the driving force for Na^+^ leak (MacMillan et al., 2015b; Olsson et al., 2016). Similar osmolyte activity occurs in some freeze-avoiding beetles (Hanzal et al., 1992). In *Drosophila*, these changes occur in response to environmental change but before a stressful cold exposure. This modification of the “resting” state is thought to limit the extent of ion balance disruption during the chilling event. The osmolytes that take the place of these extracellular ions include classical cryoprotectants like trehalose and proline, which led to the hypothesis that their accumulation may protect against ionoregulatory collapse regardless of the risk of freezing (Olsson et al., 2016). This mechanism of protection is compelling, because it would provide an evolutionary stepping-stone toward the role of cryoprotectants in SCP depression and support the notion of chill susceptibility, chill tolerance, and freeze avoidance as a continuum of thermal tolerance strategies. Indeed, chill-susceptible insects accumulate cryoprotectants in preparation for overwintering (Koštál and Šimek, 2000; Koštál et al., 2004) or when given mild cold treatments (Wang and Kang, 2005; Overgaard et al., 2007). This putative role for cryoprotectants relies on far lower concentrations of these solutes to provide protection against the cold, and prevention of ionoregulatory collapse must logically be a prerequisite for acquiring either freeze avoidance or freeze tolerance as a survival strategy. If cryoprotectants play this role in *D. ponderosae*, seasonal accumulation of glycerol and other cryoprotectants in the hemolymph may be critical to establishing a favourable equilibrium of water balance during chilling that passively protects against catastrophic hyperkalemia. By contrast, rapid production of cryoprotectants during chilling in this species (Sømme, 1964; Thompson et al., 2020), could actively prevent further progression of ionoregulatory collapse during cold stress and lead to the plateau in [K^+^] we observed here. Most likely, these means of altering osmotic gradients are combined with extensive changes to the renal system that allow for maintenance of ion homeostasis at extreme low temperatures (MacMillan et al., 2015a; Yerushalmi et al., 2018; Andersen and Overgaard, 2020). In less tolerant insects, renal plasticity plays a critical role in chill tolerance, and we expect that maintenance of ion balance in the cold is achieved by *D. ponderosae* through complementary means.

## Conclusion

In conclusion, this study unveils unique cold tolerance mechanisms in overwintering *D. ponderosae* larvae. Specifically, we show that despite possessing an impressive capacity for freeze-avoidance, these cold hardy larvae experience physiological stress and sustain injury during prolonged exposure to severe, stressful cold, which is linked to a gradual disruption of hemolymph ion balance (hyperkalemia) in a manner similar to that of less tolerant, chill-susceptible insects. The asymptotical nature of the ion balance disruption and the ability of *D. ponderosae* to stabilize hemolymph ion concentrations to just below levels causing extensive injury suggest previously unknown mechanisms of ion balance regulation and protection, which we speculate might be related to the production and accumulation of cryoprotectants. These mechanisms may stem from seasonal or rapid changes to their physiology, or a combination of both. Further research into renal and epithelial function is needed to elucidate the molecular and cellular mechanisms that promote this physiological capacity. Indeed, gaining a deeper understanding of the complex, integrative biology and physiology of overwintering in *D. ponderosae* is likely to affect future management efforts. Lastly, our findings challenge the current and historical division of cold tolerance strategies and mechanisms, and we posit that chill susceptibility, chill tolerance, and freeze avoidance represent a continuum of thermal tolerance strategies. Thus, drawing parallels between the physiological mechanisms that limit the cold tolerance of chill-susceptible insects and those that permit winter survival of more tolerant insects, like *D. ponderosae*, might promote rapid advances in our understanding of cold tolerance physiology.

## Acknowledgements

The authors would like to extend their utmost gratitude to Ashlyn Wardlaw and Dr. Anna Turbelin for their help in running experiments in the quarantine facilities at the Great Lakes Forestry Centre in Sault Ste. Marie (where both are affiliated). We would furthermore like to acknowledge Caroline Whitehouse, Glenn Dobransky, and Andrea Sharpe for their help with providing permits and guidance in collecting the beetle-infested tree logs, as well as Leanne Petro for field work associated with baiting of adult beetles to the field site and trees. Lastly, we are thankful for the assistance of Colleen Fortier, Taylor Volappi, Marion Mayerhofer, and the High Country Arborists for coordinating and assisting with the tree harvest.

## Data availability

All data has been made available to reviewers upon submission and will be linked with article on JEB’s website when published.

## Supplementary material

**Table S1.** This table contains the names and coordinates of the three weather stations used to approximate the thermal conditions of the threes containing *D. ponderosae* larvae before being transported to University of Alberta. Data was downloaded on October 18, 2023, and covered the period from September 1, 2022, to May 31, 2023.

| Weather station name | Latitude | Longitude |
| --- | --- | --- |
| Rocky Mtn House A | 52,4333 | -114,9167 |
| Edson | 53,5833 | -116,4167 |
| Edmonton Stony Plain CS | 53,5472 | -114,1084 |

**Table S2.** Sample sizes for data presented in Fig. 2, panels B-D, where supercooling points were measured at three different time points in larvae of *D. ponderosae* during overwintering.

| Season | Supercooling<br>point<br>Sample size<br>(N) |
| --- | --- |
| Autumn | 19 |
| Winter | 22 |
| Spring | 12 |

**Table S3.** Sample sizes for data presented in Fig. 3, where chill coma recovery time and survival score were quantified as a function of seasonal time point, exposure time, and exposure temperature. Note that even though the chill coma recovery time and survival score were quantified in the same animals (see Materials and Methods), the sample size for survival score is often higher than that of chill coma recovery time. This is because animals that did not recover in the 90 min time limit were counted as having not recovered and were there excluded from any analysis related to recovery times.

| Season | Exposure temperature (°C) | Exposure time (h) | Chill coma recovery time<br>Sample size (N) | Survival score<br>Sample size (N) |
| --- | --- | --- | --- | --- |
| Autumn | -10 | 0 | 11 | 11 |
| Autumn | -10 | 4 | 14 | 15 |
| Autumn | -10 | 16 | 18 | 18 |
| Autumn | -10 | 24 | 14 | 14 |
| Autumn | -10 | 48 | 12 | 13 |
| Autumn | -10 | 96 | 11 | 12 |
| Autumn | -10 | 168 | 12 | 13 |
| Winter | -20 | 0 | 15 | 15 |
| Winter | -20 | 4 | 13 | 13 |
| Winter | -20 | 16 | 12 | 14 |
| Winter | -20 | 24 | 10 | 12 |
| Winter | -20 | 48 | 15 | 16 |
| Winter | -20 | 96 | 6 | 12 |
| Winter | -20 | 168 | 9 | 12 |
| Winter | -10 | 0 | 12 | 13 |
| Winter | -10 | 4 | 16 | 16 |
| Winter | -10 | 16 | 15 | 15 |
| Winter | -10 | 24 | 16 | 16 |
| Winter | -10 | 48 | 17 | 17 |
| Winter | -10 | 96 | 11 | 12 |
| Winter | -10 | 168 | 12 | 12 |
| Spring | -10 | 0 | 13 | 13 |
| Spring | -10 | 4 | 12 | 12 |
| Spring | -10 | 16 | 16 | 16 |
| Spring | -10 | 24 | 13 | 13 |
| Spring | -10 | 48 | 11 | 11 |
| Spring | -10 | 96 | 11 | 16 |
| Spring | -10 | 168 | 18 | 18 |

**Table S4.** Sample sizes for data presented in Fig. 4, where hemolymph concentrations of K^+^ and Na^+^ were measured in overwintering larvae of *D. ponderosae* exposed to stressful low temperature for different amounts of time. Note that the sample size for Na^+^ concentration is lower than that for K^+^ concentration at the spring season for animals exposed to -10°C for 168 h because some samples were lost.

| Season | Exposure temperature (°C) | Exposure time (h) | Hemolymph K<sup>+</sup> concentration<br>Sample size (N) | Hemolymph Na<sup>+</sup> concentration<br>Sample size (N) |
| --- | --- | --- | --- | --- |
| Autumn | 22 | Room temperature control | 12 | - |
| Spring | 22 | Room temperature control | 15 | - |
| Winter | 22 | Room temperature control | 12 | - |
| Autumn | 6 | 6°C control | 9 | 9 |
| Spring | 6 | 6°C control | 15 | 15 |
| Winter | 6 | 6°C control | 7 | 7 |
| Winter | -20 | 0 | 12 | 12 |
| Autumn | -10 | 0 | 12 | 12 |
| Spring | -10 | 0 | 16 | 16 |
| Winter | -10 | 0 | 10 | 10 |
| Winter | -20 | 4 | 13 | 13 |
| Autumn | -10 | 4 | 14 | 14 |
| Spring | -10 | 4 | 10 | 10 |
| Winter | -10 | 4 | 12 | 12 |
| Winter | -20 | 16 | 15 | - |
| Autumn | -10 | 16 | 15 | - |
| Spring | -10 | 16 | 16 | - |
| Winter | -10 | 16 | 12 | - |
| Winter | -20 | 24 | 10 | 10 |
| Autumn | -10 | 24 | 13 | 13 |
| Spring | -10 | 24 | 15 | 15 |
| Winter | -10 | 24 | 9 | 9 |
| Winter | -20 | 48 | 16 | - |
| Autumn | -10 | 48 | 11 | - |
| Spring | -10 | 48 | 9 | - |
| Winter | -10 | 48 | 17 | - |
| Winter | -20 | 96 | 6 | 6 |
| Autumn | -10 | 96 | 12 | 12 |
| Spring | -10 | 96 | 16 | 16 |
| Winter | -10 | 96 | 7 | 7 |
| Winter | -20 | 168 | 9 | 9 |
| Autumn | -10 | 168 | 11 | 11 |
| Spring | -10 | 168 | 17 | 12 |
| Winter | -10 | 168 | 8 | 8 |

**Fig. S1.**
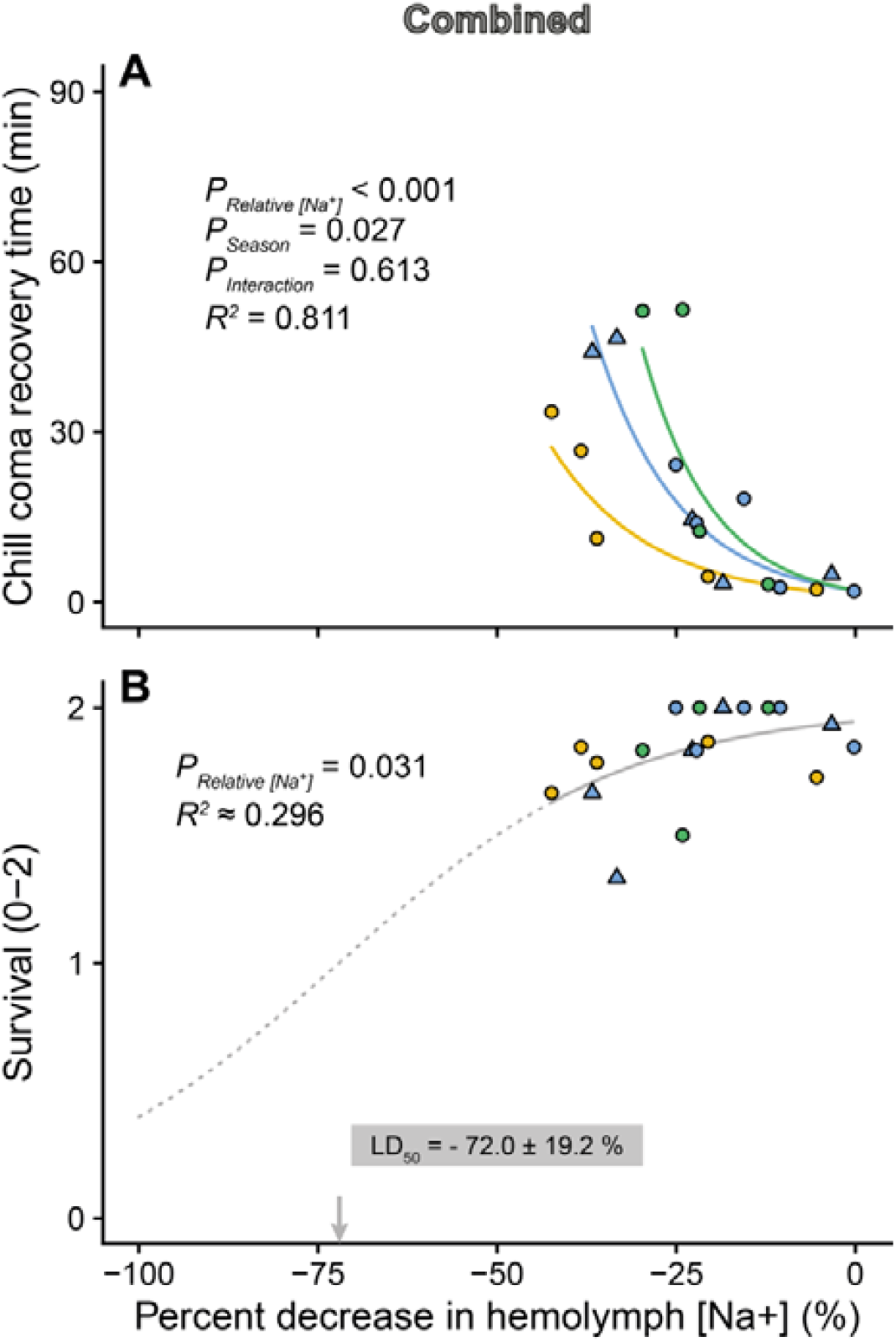
Correlations between the degree of hemolymph hyponatremia and cold tolerance phenotypes. These correlations were made using measurements of CCRT, survival score, and hemolymph Na^+^ concentration obtained at the same time points from autumn (yellow), winter (blue), and spring (green), and at both -10°C (circles) and -20°C (which was only measured in winter). Note that these measurements are from a subsample of time points. Overall there was a strong exponential relationship between the degree of hemolymph hypernatremia and the CCRT (**A**), and this relationship was season-specific such that recovery times were shortest in the fall, followed by winter and then spring (no effect of exposure temperature were found). In terms of the survival score (B), there were no effects of temperature or seasonality and survival was found to decrease as a function hyponatremia. Like with hemolymph hyperkalemia, we fitted a sigmoidal model to the data which we used to extrapolate and find a LD50 at a -72.0 ± 19.2% decrease in hemolymph Na^+^ concentration.

